# Islet amyloid disrupts MHC Class II antigen presentation and protects NOD mice from autoimmune diabetes

**DOI:** 10.1101/2024.09.10.612362

**Authors:** Heather C Denroche, Tori Ng, Jane Velghe, Imelda Suen, Liam Stanley, Dominika Nackiewicz, Mitsu Komba, Sam Chen, Galina Soukhatcheva, Lei Dai, C. Bruce Verchere

**Affiliations:** Canucks for Kids Fund Childhood Diabetes Laboratories, BC Children’s Hospital Research Institute, Departments of Surgery, Centre for Molecular Medicine and Therapeutics, University of British Columbia, Vancouver, British Columbia, Canada; Pathology and Laboratory Medicine, Centre for Molecular Medicine and Therapeutics, University of British Columbia, Vancouver, British Columbia, Canada; Integrated Nanotherapeutics Inc., Burnaby, Canada

## Abstract

Islet amyloid contributes to beta cell failure in type 2 diabetes through several mechanisms, one being the potent induction of local islet inflammation through activating inflammatory pathways in islet macrophages. We performed an unbiased phenotypic investigation of islet macrophages in the early stage of islet amyloid formation using single cell RNA sequencing of resident islet macrophages in mice with and without the amyloidogenic form of human islet amyloid polypeptide (hIAPP). This revealed that MHC Class II antigen presentation genes were strongly down-regulated in islet macrophages during islet amyloid formation. As islet amyloid has recently been reported in pancreases of people with type 1 diabetes, we sought to investigate the impact of islet amyloid in the NOD mouse model of type 1 diabetes. Both overexpression and physiological expression of hIAPP delayed diabetes in NOD mice relative to littermate controls, corresponding with decreased markers of antigen presentation and activation, as well as decreased immune cell infiltration in islets. Adoptive transfer studies showed that systemic autoimmune function remained intact and beta cells from hIAPP transgenic mice did not evade immune recognition by diabetogenic T cells, collectively indicating the protection from diabetes was mediated by localized disruption of antigen presentation in the pancreas. Consistent with this, incubation of dendritic cells with IAPP aggregates decreased MHC Class II surface expression and diminished antigen-specific T cell activation in vitro, through a phagocytosis-dependent mechanism.

Collectively our data show that despite the well-established pro-inflammatory response of macrophages to IAPP aggregates, the uptake of IAPP aggregates during early amyloid formation also disrupts MHC Class II antigen presentation and slows beta cell autoimmunity.

## Introduction

Type 1 diabetes is caused by autoreactive T cell-mediated destruction of pancreatic beta cells leading to insufficient insulin levels and consequently hyperglycemia. Islet macrophages play a key role in the initiation and progression of type 1 diabetes pathogenesis and depletion of macrophages protects NOD mice from developing diabetes [1–4]. The secretion of pro-inflammatory cytokines by macrophages contributes to beta cell loss in type 1 diabetes, exemplified by the protective effects of deleting NLRP3 inflammasome in NOD mice and inhibiting TNF in humans with recent onset type 1 diabetes [5–7]. In addition, islet macrophages are also important antigen presenting cells (APCs), phagocytosing secreted peptides, insulin granules and crinosomes from beta cells and presenting beta cell antigens to autoreactive T cells [1–3, 8–10].

Islet amyloid, formed by the aggregation of the beta cell peptide hormone islet amyloid polypeptide (IAPP), is a strong proinflammatory stimuli, triggering islet inflammation contributing to beta cell dysfunction [11–15]. Islet macrophages, residing in close contact with beta cells and sampling insulin granule content and secretions [10, 16] are poised to directly interact with IAPP aggregates as they form. We and others have shown in vitro that IAPP aggregates directly interact with TLR2 and are phagocytosed by macrophages, eliciting a pro-inflammatory response and NLRP3 inflammasome activation [15, 17, 18]. Furthermore, by systemically depleting macrophages or inhibiting inflammatory cytokine signalling, we have shown macrophage inflammatory responses are an essential link between IAPP aggregates and beta cell dysfunction [11–13]. Nevertheless, the direct impact of islet amyloid on resident islet macrophages in situ has not been studied.

In this study, we investigated the effects of islet amyloid on islet macrophages in situ in an unbiased manner by performing single cell RNA sequencing on islet macrophages from a mouse model of early islet amyloid formation (hIAPP Tg/0 mice). This revealed a surprising result that MHC Class II antigen presentation genes are significantly down-regulated in islet macrophages in the presence of islet amyloid. While islet amyloid has long been considered a pathology unique to type 2 diabetes, it has recently been found in islets from people with type 1 diabetes [19–21]. Consequently, we investigated the effect of IAPP aggregates on antigen presentation and the role of islet amyloid in autoimmune-mediated diabetes in the NOD mouse model.

## Methods

### Animals

All animal experiments were approved by the Animal Care Committee at the University of British

Columbia and performed in agreement with the ethical guidelines and policies provided by the Canadian Council on Animal Care. All animal experiments were conducted at BC Children’s Hospital Research Institute where mice were housed under 12-h light 12-h dark conditions at room temperature, with ad libitum access to chow (Teklad, 2918, Envigo, Indianapolis, IN, US; Calorie distribution: 24% protein, 18% fat, 58% carbohydrate) and water. Where indicated, mice were placed on high fat diet (HFD; D12451, Research Diets, New Brunswick, NJ, USA: Calorie distribution: 20% protein, 45% fat, 35% carbohydrate) for 2 weeks.

C57BL/6J (000664), NOD/ShiLtJ (NOD, 001976), NOD.Cg-Prkdcscid/J (NOD SCID, 001303), B6.Cg-Tg(TcraTcrb)425Cbn/J (OT-II, 004194), FVB/N-Tg(Ins2-IAPP)RHFSoel/J (hIAPP Tg/0, RIPHAT, 008232), BALB/cJ (000651) mice were purchased from Jackson Laboratory, Bar Harbor, Maine, USA. hIAPP knockin (NOD.Iapp^h/h^) mice were developed by Hiddinga et al. [22] obtained from Dr. Steven Kahn (University of Washington Diabetes Institute, Seattle, USA).

### Randomisation

All animals were assigned to their respective groups depending on genotypes and littermate controls were used for all in vivo experiments.

### Single cell RNA sequencing

hIAPP Tg/0 and hIAPP 0/0 littermates on a mixed C57BL/6J x FVB/NJ background, were generated by crossing hIAPP Tg/0 transgenic mice on the FVB/NJ background with C75BL/6J mice. After HFD, islets were isolated and pooled (5 mice per sample, n=3 samples per genotype) for FACS enrichment of islet immune cells via BD FACS ARIA II (BD Biosciences, San Jose, CA, USA) into PBS containing 2% FBS (Thermo Fisher Scientific, Canada), pH 7.4. Based on the total amount of cells sorted into the CD45+ fraction, the amount of CD45- cells to spike back in was calculated to achieve a final ratio of 70% CD45+ cells to 30% CD45- cells. Each combined sample was loaded onto the Chromium Controller and underwent library preparation single-cell RNA sequencing using the Chromium Single Cell 3’ V2 Reagent Kit (10X Genomics). Sample libraries were then sequenced on an Illumina NextSeq500 to a read-depth minimum of 80,000 reads per cell. Reads were aligned to a reference genome using CellRanger v2.1.1 (10X Genomics). Resulting sample gene expression matrices were then each individually processed in R.

### Transcriptomic analysis

Sample gene expression matrices were each individually processed in R using various packages before downstream analysis to remove confounding variables. First, the effect of ambient mRNA detected in droplets was mitigated using SoupX (1.6.2) [23]. Next, gene expression matrices were imported into RStudio using Seurat (4.3.0) [24]. For each sample, low quality cells were removed prior to applying DoubletFinder (2.0.4), to detect and remove artificial multiplets before performing SCTransform, selecting 3000 integration features, and integrating samples into one dataset [24][25][26]. Next, a standard workflow was run for visualization and clustering: principle component analysis (Seurat, RunPCA, npcs=30), dimensionality reduction (Seurat, RunUMAP, reduction = "pca", dims = 1:30), construction of nearest neighbour graph (Seurat, FindNeighbors, reduction = "pca", dims=1:30) and cluster detection (Seurat, FindClusters, res=0.8, algorithm=1) [24]. Cell types were annotated based on known marker genes. Gene counts of all the cells from the same sample for each cell type were summed (Seurat, AggregateExpression) to generate psudobulk gene expression profiles [27]. Differentially expressed genes (DEGs) between the transgenic and wild type conditions for each type were calculated on the pseudobulk dataset (Seurat, FindMarkers) using the DESeq2 test (1.36.0) [24][28]. Gene set enrichment analysis (GSEA) was performed using fGSEA (1.31.3) [29].

### Islet isolation

Islets were isolated via collagenase injection into the pancreatic duct, and dispersed for flow cytometry as previously published [30, 31]. Briefly, islets were harvested from mice and hand-picked to purity in complete RPMI 1640 media (Thermo Fisher Scientific Canada) and subsequently exocrine-free islets were selected and dispersed with 0.02 % trypsin for 3 min at 37 °C and washed with FACS buffer (2% FBS, 2 mM EDTA (Thermo Fisher Scientific) in PBS (Thermo Fisher Scientific), pH7.4) for downstream staining and analysis.

### Splenocyte and lymph node isolation

Spleens and lymph nodes were harvested from mice and homogenized through a 40 µm cell strainer. Cells were washed once with PBS, pH7.4 (Thermo Fisher Scientific), then lysed in ammonium chloride solution (Stemcell Techonologies, Canada) at 4 °C for 10 minutes. Following lysis, cells were washed once with PBS and resuspended in PBS for counting and flow cytometry.

### Flow cytometry

Cells were incubated in Fc block (Table 1) for 10 min prior to addition of antibodies. Antibodies and staining reagents are provided in 1. Fixable viability eFlour780 (Thermo Fisher Scientific) or 7-AAD (Thermo Fisher Scientific) were used according to manufacturer instructions to measure viability. Flow cytometry was conducted on a LSR Fortessa (BD Biosciences, USA), Cytoflex (Beckman Coulter, USA) or cells were sorted by FACS as above. Data were analyzed by FlowJo software (BD Biosciences, USA).

**Table 1.** Flow cytometry reagents and antibodies.

| *Marker* | *Clone* | *Dilution* |
| --- | --- | --- |
| CD3 | 17a2 | 1:100 |
| CD4 | RM4-5 | 1:100 |
| CD8 | 53-6.7 | 1:100 |
| CD11b | M1/70 | 1:100 |
| CD11c | N418 | 1:100 |
| CD19 | eBio1D3 | 1:100 |
| CD45 | 30-F11 | 1:100 |
| Fc block (CD16/CD32) | 93 | 1:100 |
| Foxp3 | FJK-16s | 1:25 |
| FVD 780 | n/a | 1:1000 |
| MHC II I-A/d | 39-10-08 | 1:100 |
| MHC II I-A/I-E | M5/114.15.2 | 1:100 |

### NOD diabetes progression studies

hIAPP Tg/0 transgenic mice on the FVB/NJ background were backcrossed (>10 generations) with NOD mice to generate NOD.hIAPP Tg/0 and NOD.hIAPP 0/0 littermates. Iapp^h/h^knockin mice on a C57BL/6J background [22] were backcrossed (>10 generations) with NOD mice to create NOD.Iapp^h/m^ mice.

NOD.Iapp^h/h^, NOD.Iapp^h/m^ and NOD.Iapp^m/m^ littermates were generated via NOD.Iapp^h/m^ heterozygous breeding. Blood glucose and body weight were monitored 1-2x weekly for up to 30 weeks of age.

Diabetes onset was defined as 2 consecutive blood glucose measures ≥ 20 mM.

### Genotyping, primers

All hIAPP Tg/0 strains described in this paper were maintained as hemizygous carriers, by crossing male carriers to wildtype females. All mice were genotyped by DNA extracted from ear notches following standard PCR protocols. The hIAPP transgene was genotyped with the primers listed in Table 2. Knock-in hIAPP mice were genotyped with the primers listed in Table 2. For generation of NOD SCID.hIAPP Tg/0 mice, the Prkdc^scid^ allele was genotyped using restriction fragment length polymorphism assay described Quadros et al. [32].

**Table 2.**
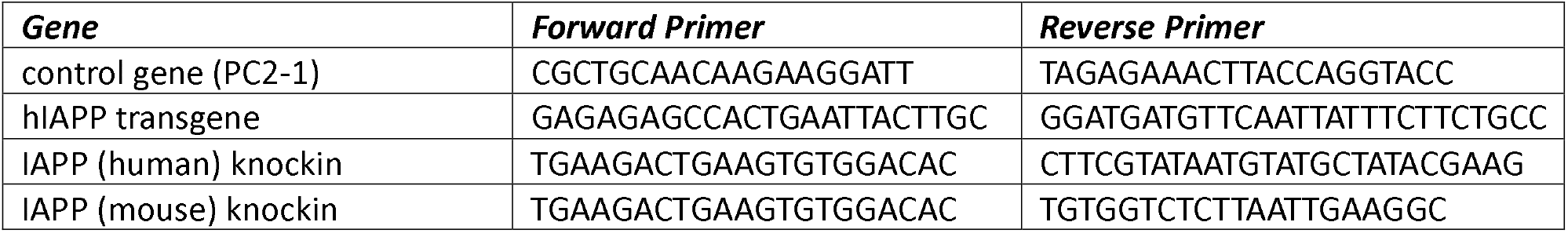
Primers and sequences.

### Adoptive transfers

For a subset of the adoptive transfer experiments, recipient NOD SCID.hIAPP Tg/0 and NOD SCID.hIAPP 0/0 littermates were generated by backcrossing the NOD.hIAPP Tg/0 transgenic mice with homozygous NOD.Cg-Prkdc^scid/scid^ mice to generate 50% NOD.Prkdc^scid/wt^ ; hIAPP Tg/0 mice which were then crossed again to NOD SCID mice to generate NOD.Prkdc^scid/scid^; hIAPP Tg/0 and NOD.Prkdc^scid/scid^; hIAPP 0/0 mice which were then intercrossed to maintain homozygosity for the Prkdc^scid^ allele. Splenocytes from 18- week old female NOD/ShiLtJ mice were isolated and injected at a dose of 10 million cells per recipient into the tail vein of 17-week old female NOD SCID.hIAPP Tg/0 and NOD SCID.hIAPP 0/0 female mice. In another experiment, splenocytes from non-diabetic 19-week old female NOD.hIAPP Tg/0 and NOD.hIAPP 0/0 mice were injected at a dose of 10 million splenocytes per recipient into the tail vein of 25-week old female NOD SCID mice. Blood glucose was measured 2x per week in recipients and diabetes defined as 2x blood glucose >13 mM.

### Allo-transplantation studies

Diabetes was induced in 8-week old Balb/cJ mice by injection of 200 mg/kg streptozotocin (#S0130, Sigma, US) prepared as previously described [33]. Islets from 25-week old hIAPP Tg/0 and hIAPP 0/0 male mice (to ensure amyloid formation) on the FVB/NJ background were isolated and transplanted under the kidney capsule of sex-matched diabetic Balb/cJ recipient mice following previously published protocols [34–36]. Diabetes was defined as 2x blood glucose > 20 mM.

### Metabolic Tests

For glucose tolerance tests, all mice were fasted for 6 h and subsequently injected intraperitoneally with glucose dissolved in PBS (dose is indicated in the figure legend). Tail blood glucose was measured using a OneTouch Ultra Glucometer (Life Scan Inc., Burnaby, BC, Canada) at indicated time points post injection. To measure in vivo glucose-stimulated insulin levels (GSIL), blood was collected from the saphenous vein after a 6-h fast and after injection of glucose for the measurement of plasma insulin via Stellux Chemiluminescent Rodent Insulin ELISA (#80-INSM R-CH01, Alpco, Salem, NH, USA). Ex vivo glucose-stimulated insulin secretion (GSIS) was measured as previously described [31].

### T cell proliferation assay

Synthetic hIAPP and rIAPP (Bachem, Switzerland) were prepared as previously published [17]. Bone marrow-derived dendritic cells (BMDCs) were generated from C57BL/6J mice, and treated with synthetic hIAPP, rIAPP (10 µM) or vehicle as indicated for 24 h, with or without 10 ng/ml LPS (Sigma).

Subsequently, BMDCs were washed and stained for flow cytometry or co-cultured at a 1:5 ratio with CFDA-SE (Stemcell Technologies, Canada) -labelled CD4 T cells isolated from a B6. Cg-Tg(TcraTcrb)425Cbn/J (OT-II) donor mouse in RPMI-1640, (Thermo Fisher Scientific) supplemented with 10% heat inactivated FBS (Thermo Fisher Scientific, Burnaby, BC, Canada), 100U penicillin/streptomycin (Thermo Fisher Scientific), 10 mM HEPES (Thermo Fisher Scientific), 50 μM β-mercaptoethanol (MilliporeSigma, Canada), 1x Glutamax (Thermo Fisher Scientific, Canada), and 10mM sodium pyruvate (Thermo Fisher Scientific). After 3 days, cells were stained for flow cytometry as above with CD4 antibody (Table 1), fixable viability dye eFlour780 (Thermo Fisher, US) and analyzed by flow cytometry on a Cytoflex. Proliferation was determined by CFSE dilution as a percent CD4 cells, after gating on viable singlets.

### Nanostring

Islet gene expression in NOD.hIAPP Tg/0 mice and littermate controls was performed via Nanostring nCounter XT (Nanostring Technologies, USA) with a custom Code Set. Cell lysates were prepared and added to hybridization mixtures for 18 h at 65°C with a 70°C-heated lid according to manufacturer instructions. Gene expression was measured with a Nanostring SPRINT Profiler, and data were analyzed via Nsolver 4.0 software.

### Immunization

Ovalbumin (Ova) mRNA with 5’ methoxy-uridine (5moU) base modification (Trilink BioTechnologies, USA) was formulated into lipid nanoparticles by Integrated Nanotherapeutics (Canada) with their proprietary ionizable lipid using a rapid mixing process [37]. Briefly, all lipid components, were dissolved in ethanol while the mRNA was dissolved in pH 4.0 buffer. The ethanol and buffer solutions were rapidly mixed at a ratio of 1:3, respectively. The resultant LNP solutions were dialyzed overnight in pH 7.4 buffer to remove residual ethanol and to increase the pH to physiological levels. All LNP-mRNA were formulated at an amine-to-phosphate ratio (N/P) of 6. Lipid concentrations were determined by measuring cholesterol content (Cholesterol Total E Assay, Wako Chemicals, Richmond, VA, USA) and RNA entrapment and concentrations were determined using Ribogreen (Thermo Fisher Scientific, Burnaby, BC, Canada). Final LNP-mRNA were stored frozen at -70°C. Mice were injected with 10 µg mRNA in LNP formulation ip at 0 and 14 days, prior to collection of blood from the saphenous vein for measurement of anti-ova IgG1 by Cayman Chemical ELISA (Cayman Chemical, USA).

### Immunohistochemistry staining

Pancreases were harvested immediately after euthanasia, and processed and stained as previously described [38] then fixed with 4% paraformaldehyde for 24h. Samples were washed with 70% ethanol prior to paraffin embedding and sectioning (5 μm). Following removal of paraffin layer and rehydration, samples were washed with PBS (Thermo Fisher Scientific) and blocked for 1h at room temperature (Dako Protein block, Agilent Technologies, USA). Samples were incubated overnight at 4°C in anti-insulin antibody (1:500 Dako, Agilent Technologies), washed in 3 times in PBS and incubated for 1 h at room temperature in Alexa 594 goat anti-guinea pig secondary antibody (1:200, Thermo Fisher Scientific) in a dark humid chamber, followed by incubation in 0.5% (wt/vol) thioflavin S (Sigma) for 2 min, and washed with 70% ethanol and water prior to mounting. Images were acquired on a BX61 microscope (Olympus, Richmond Hill, ON, Canada) using 10× air objective.

## Statistical methods

With the exception of gene expression data, all statistical analyses were performed using GraphPad Prism 10.0 (GraphPad Software, La Jolla, CA, USA). Data are presented as mean ± SEM, with individual data points from biological replicates, unless otherwise indicated. For comparisons of only two groups, normally distributed data were analyzed by t-test with Welch’s correction whereas non-normal data were analyzed by Mann-Whitney test. Data were tested for normality by the Shapiro-Wilk test. For comparisons of more than two groups, normally distributed data were analyzed by one-way ANOVA and non-normal data analyzed by Kruskal-Wallis test. For repeated measures of two or more groups data were analyzed by two-way repeated measures ANOVA with Geisser-Greenhouse correction for non-sphericity. Diabetes incidence data were analysed via logrank test. The number of observations (n) are provided in figure legends.

## Results

### The MHC Class II pathway is downregulated in islet macrophages of hIAPP Tg/0 mice

We first sought to examine the phenotype of islet macrophages exposed to islet amyloid in situ.

Unlike human IAPP, rodent IAPP does not form amyloid due to differences in its primary amino acid sequence, and thus rodents do not normally develop islet amyloid. Therefore, we used mice carrying a transgene expressing the amyloidogenic, human form of IAPP under control of the rat insulin 2 promoter (hIAPP Tg) which drives IAPP overexpression in beta cells. Under obesogenic conditions, hIAPP Tg/0 mice overexpress hIAPP in beta cells and develop islet amyloid and beta cell dysfunction modelling type 2 diabetes [12–14, 39, 40]. hIAPP Tg/0 mice on a FVB/NJ background were crossed with C75BL/6J mice to generate hIAPP Tg/0 and hIAPP 0/0 littermates on a F1 C57BL6/J x FVB/NJ background (Fig 1A) which helps promote islet amyloid and beta cell dysfunction on HFD [39]. To induce early islet amyloid formation, 8-week old male hIAPP Tg/0 and hIAPP 0/0 male mice, which have similar baseline glucose tolerance at this age (Fig 1B) were placed on HFD for 2 weeks. Following this brief dietary change, glucose tolerance (Fig 1C) and fasting blood glucose (Fig 1D) worsened in hIAPP Tg/0 mice relative to littermate hIAPP 0/0 controls despite maintaining similar body weight (Fig 1E), indicative of early islet amyloid formation disrupting beta cell function.

**Figure 1.**
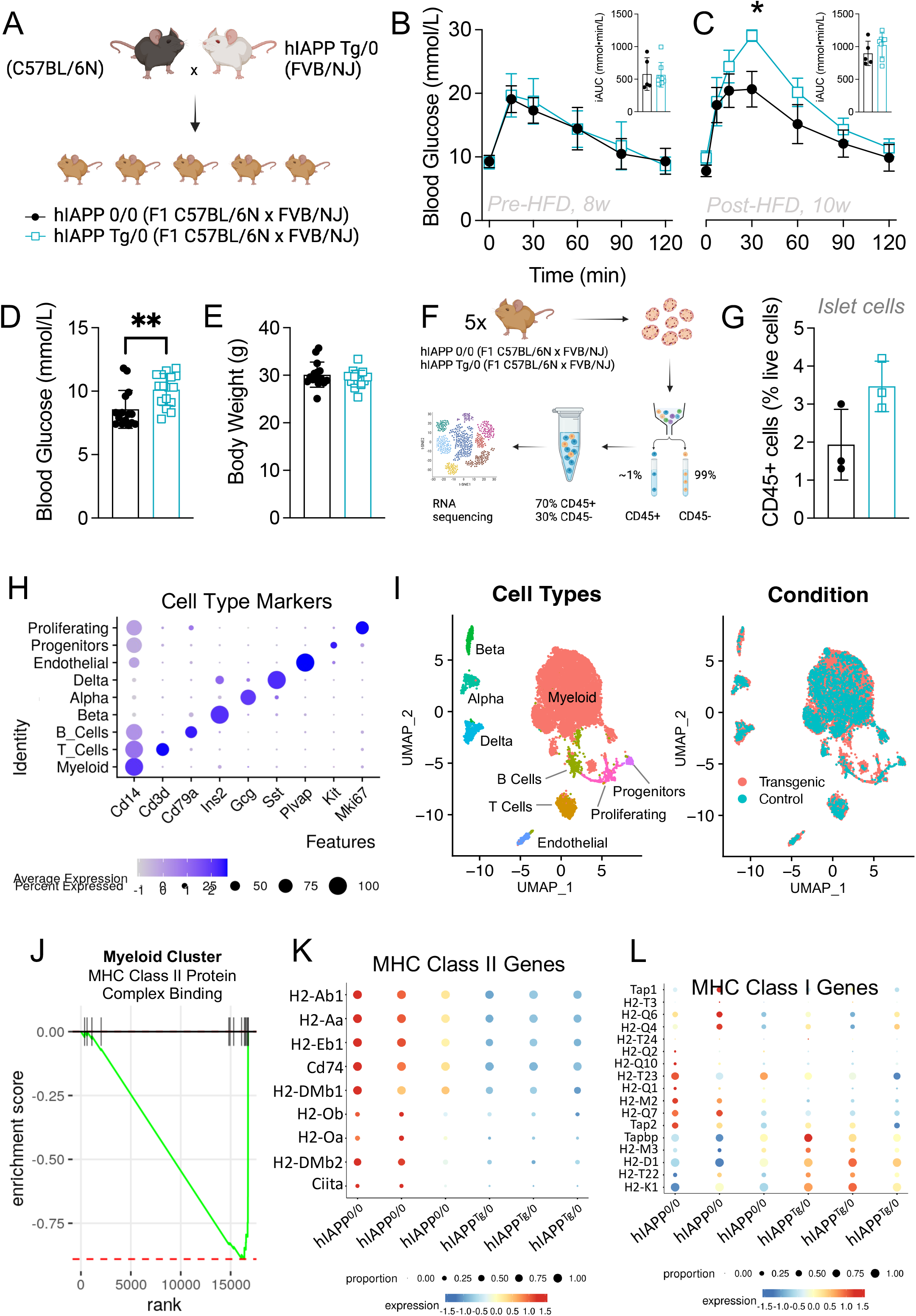
Downregulation of MHC Class II pathway in islet macrophages of hIAPP Tg/0 mice. A-G) Data from female control (black, solid circle) and hIAPP Tg/0 (teal, open square) transgenic mice on a F1 C57BL/6N x FVB/NJ background. A) Overview of animal model breeding strategy (made in BioRender). Wildtype C57BL/6N mice were crossed with hIAPP Tg/0 transgenic FVB/NJ mice to generate F1 littermate hIAPP Tg/0 transgenic hIAPP 0/0 control mice on a mixed C57BL/6N x FVB/NJ background. B) Glucose Tolerance data (1.5g glucose/kg BW, i.p) from chow-fed 8-week-old and. C) 10-week old (Following 2 weeks of HFD feeding) mice (n (hIAPP 0/0): week 8 = 5, week 10 = 5), (hIAPP Tg/0): week 8 = 5, week 10 = 5). Data were analysed using repeated measures mixed effects analysis. Individual incremental AUCs from the experimental and control groups were analysed via Mann-Whitney tests. D) 4h fasted blood glucose levels (mM) and E) Body weights (g) of 10-week old mice fed a HFD for 2 weeks (n(hIAPP 0/0)=15; n(hIAPP 0/0)=16). BG and BW data were analyzed using Mann-Whiteny and unpaired t-tests, respectively, depending on normality distribution F) Overview of single cell RNA seq strategy islet CD45+ cells (made in BioRender). G) % positive CD45+ cells of total live islet cells (hIAPP Tg/0: n=3; hIAPP 0/0: n=3). H) Dot plot showing key marker gene expression in each major scRNAseq population. I) UMAP plots of major scRNAseq populations. J) Enrichment score of GO Molecular Function MHC Class II Protein Complex Binding gene set from hIAPP Tg/0 myeloid cell population vs. control. K) Dot plot of select MHC Class II associated genes across samples. L) Dot plot of select MHC Class I associated genes across samples. Data are shown as mean +/- SD. p < 0.05 (*), p < 0.01 (**),p < 0.0001 (****).

Next, we used single cell RNA sequencing to examine the islet macrophage response during early islet amyloid exposure in an unbiased manner. Islets were isolated and dispersed to single cell suspensions from hIAPP Tg/0 and hIAPP 0/0 mice (Fig 1F). To obtain sufficient islet immune cells (representing only ∼2% of total islet cells), islet cell suspensions from 5 mice were pooled per sample, and viable CD45+ cells sorted by FACS. We observed increased CD45+ cell frequency in dispersed islet cells from hIAPP Tg/0 mice relative to hIAPP 0/0 mice (Fig 1G) consistent with islet inflammation [11, 13, 14]. This finding was not significant after only 2 weeks of HFD, with a sample number of 3 per group, where islets from 5 mice were pooled per sample. Each sorted CD45+ sample was spiked with CD45- islet cells to reconstitute samples of 75% CD45+ 25% CD45- islet cells for scRNAseq (Fig 1F). scRNAseq confirmed this composition; samples consisted of a small proportion of endocrine cells (high ChgA expression, Supplemental Figure 1) with distinct beta cell, delta cell and alpha cell clusters (Figure 1H-I), along with a large population of immune cells (marked by Ptprc expression, Supplemental Fig 1) and a small population of endothelial cells (marked by Plvap). Myeloid cells comprised the majority of islet immune cells (high expression of Cd14 and low expression of Cd3e, Cd79a), with small subsets of T and B cells (high Cd3e, Cd79a respectively). These proportions amongst the immune cell population are consistent with previous work confirming 90% islet immune cells are islet macrophages and approximately 2-5% of immune cells are T and B cells [30, 31, 41].

Though islet amyloid is known to trigger inflammation in macrophages, to our surprise MHC Class II pathway was strongly down-regulated in the myeloid population of hIAPP Tg/0 islets compared to wildtype controls (Figure 1J). Interestingly, transcripts of both MHC Class II complexes and genes involved in MHC Class II antigen processing, for example Cd74, were similarly down-regulated in hIAPP Tg/0 islet macrophages (Figure 1K). In addition, Ciita, the master transcriptional regulator of MHC Class II genes, was also down-regulated (Figure 1K). In contrast, MHC Class I genes were not significantly altered in hIAPP Tg/0 islet macrophages despite being proximal to MHC Class II genes in the H2 locus on chromosome 17 (Figure 1L). Collectively, these data suggest that despite its known inflammatory effects on islet macrophages [11, 13–15], one of the strongest effects of early islet amyloid exposure on islet macrophages is a specific down-regulation of MHC Class II antigen presentation pathway.

### Islet amyloid delays autoimmune diabetes in NOD mice

Given the surprising finding that MHC Class II antigen presentation in islet macrophages is down- regulated in early islet amyloid formation, and since islet amyloid has recently been identified in pancreases of people with type 1 diabetes [19–21], we next sought to investigate the effect of islet amyloid in the NOD mouse model of autoimmune diabetes. Islet amyloid has been studied extensively in mouse models of type 2 diabetes and syngeneic islet transplantation using hIAPP expressing mouse models, but has yet to be studied in mouse models of type 1 diabetes. Thus, we generated a novel model of islet amyloid formation in type 1 diabetes, by backcrossing hIAPP Tg/0 FVB/NJ mice > 10 times to the NOD/ShiLtJ background to generate NOD.hIAPP Tg/0 and control NOD.hIAPP 0/0 littermates (Fig 2A). Interestingly, NOD.hIAPP Tg/0 mice had a significantly reduced incidence of spontaneous autoimmune diabetes relative to their littermate controls (Fig 2A, p = 0.016). By 30 weeks of age, diabetes incidence was 55.5% for NOD.hIAPP Tg/0 mice with a median onset of 30.3 weeks in comparison to a diabetes incidence of 73.3% and a median onset of 19.5 weeks for NOD.hIAPP 0/0 control mice. In males, the median onset and diabetes incidence appeared to be similar between NOD.hIAPP Tg/0 (27.6, 58%) and littermate control (27.5, 54%) mice (Supplemental Figure 2).

**Figure 2.**
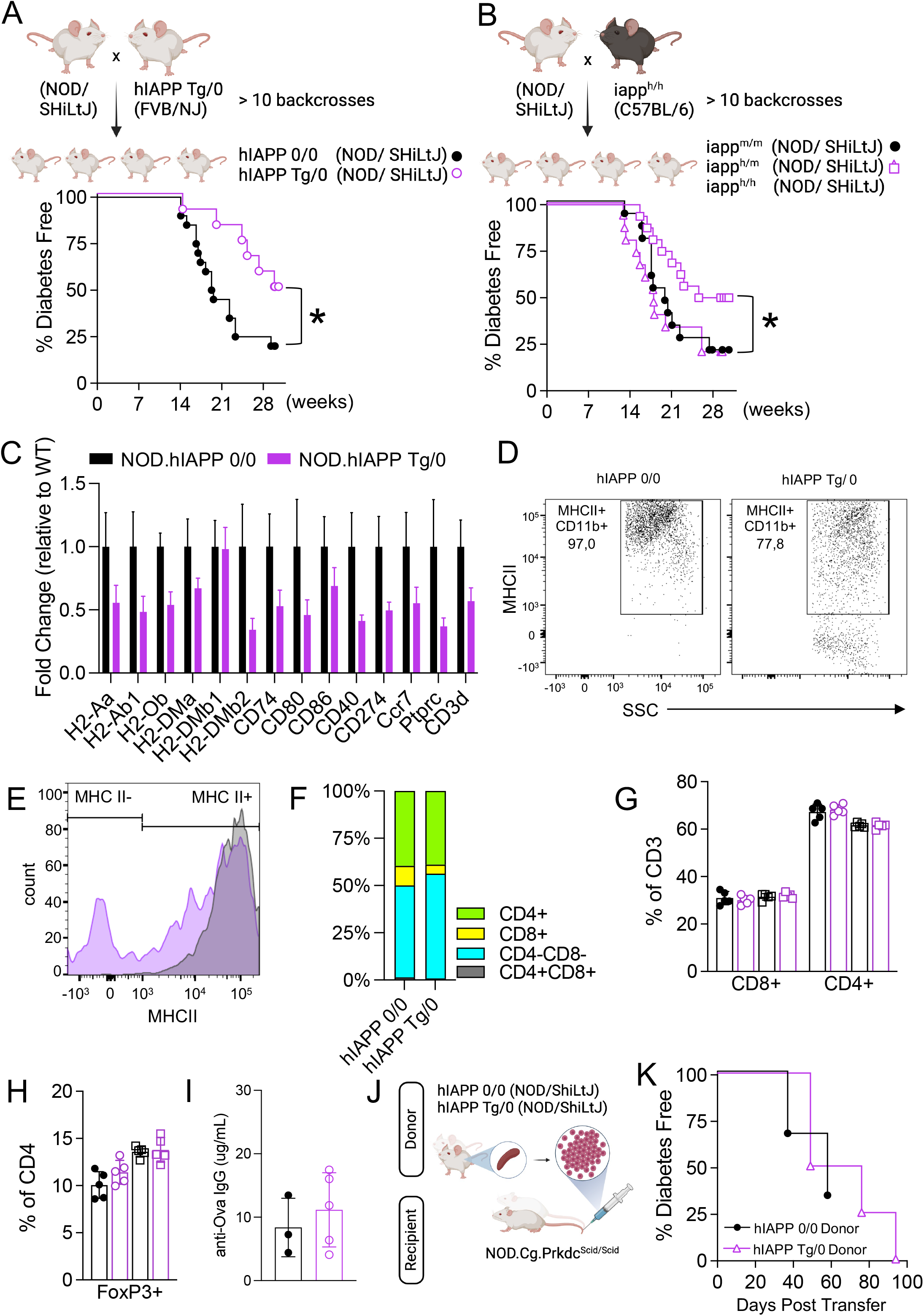
human IAPP delays spontaneous diabetes in female NOD mouse models. A-G) Data from female control (black, solid circle) and hIAPP Tg/0 (pink, open circle) transgenic mice on a NOD SHiLtJ background. A) Overview of hIAPP transgenic NOD/SHiLtJ animal model breeding strategy (made in BioRender) and diabetes incidence. hIAPP Tg/0 mice were fully backcrossed from a FVB/NJ background to a NOD SHiLtJ background (> 10 backcrosses). Diabetes incidence of female NOD.hIAPP Tg/0 (n= 9) and NOD.hIAPP 0/0 control (n=14) mice. Data were analyzed via a logrank test. B) Overview of hIAPP knockin NOD/SHiLtJ animal model breeding strategy (made in BioRender) and diabetes incidence. NOD.Iapp^h/h^ C57BL/6-J mice on a were fully backcrossed to an NOD SHiLtJ background (> 10 backcrosses). Diabetes incidence of NOD.Iapp^m/m^ (black, solid circle, n=10), NOD.Iapp^m/h^ (pink, open triangle, n=8), and NOD.Iapp^h/h^ (pink, open triangle, n=10) mice. Data were analyzed via logrank test. C) Nanostring gene expression data from islets isolated from female NOD.hIAPP 0/0 (black) and NOD.hIAPP Tg/0 (pink) mice n=6/group, data are mean + SEM of fold change relative to NOD.hIAPP 0/0. D) Representative FACS plot of islet macrophages from NOD.hIAPP 0/0 and NOD.hIAPP Tg/0 mice. E) Modal normalized count vs MHC II fluorescence in islet macrophages. F) Islet CD4+ CD8+ islet infiltration data from female NOD.hIAPP Tg/0 and NOD.hIAPP 0/0 mice. G) CD4+, CD8+ T cell frequency in spleen (circular symbols) or the pancreatic lymph node (square symbols). H) Treg cell frequency in spleen (circular symbols) or the pancreatic lymph node (square symbols). I) Serum Ova IgG1 concentrations from NOD.hIAPP Tg/0 and NOD.hIAPP 0/0 mice immunized with mRNA vaccines encoding Ovalbumin. J) Schematic overview of adoptive transfer of prediabetic splenocytes from either NOD.hIAPP Tg/0 or NOD.hIAPP 0/0 donor mice into NOD.CG.Prkdc^Scid/Scid^ recipient mice (made in BioRender). K) Diabetes incidence curve of NOD.CG.Prkdc^Scid/Scid^ recipient mice that have received prediabetic splenocytes from NOD.hIAPp 0/0 (black solid circular symbol, n=3) or NOD.hIAPP Tg/0 (pink open triangular symbol, n=4) donor mice. Data were analyzed via logrank test. Data are shown as mean +/- SD. p < 0.05 (*), p < 0.01 (**),p < 0.0001 (****).

To control for artifacts that could result from the hIAPP transgene, we generated a complimentary NOD/ShiLtJ mouse model carrying hIAPP protein coding sequence knocked-in to the endogenous mouse Iapp locus. hIAPP knock-in C57BL/6 mice [22] were backcrossed to NOD/ShiLtJ mice for > 10 generations to generate littermates homozygous for either human IAPP (NOD.Iapp^h/h^), wildtype mouse IAPP (NOD.Iapp^m/m^) and heterozygous (NOD.Iapp^h/m^) (Fig 2B). Similar to the results of the transgenic NOD.hIAPP mice, homozygous hIAPP knock-in NOD mice were protected from diabetes relative to homozygous wildtype controls (median onset 28.2 vs 18.0 and total incidence 50% vs 80%, respectively) (Fig 2B). Interestingly, heterozygous hIAPP knock-in NOD.Iapp^h/m^ mice were not protected from diabetes (median onset 19.9 and total incidence 85%) (Fig 2B). In summary, these data show that in two distinct NOD mouse models of islet amyloid formation, hIAPP delays autoimmune diabetes progression, with protection possibly dependent on the level of hIAPP expression.

We next examined the immune phenotype of NOD.hIAPP Tg/0 mice. In agreement with an effect on antigen presentation, whole islets isolated from NOD.hIAPP Tg/0 mice had reduced MHC Class II antigen presentation gene expression and APC activation relative to littermate controls (Fig 2C).

Furthermore, macrophages isolated from NOD.hIAPP Tg/0 islets had decreased mean fluorescence intensity of MHC II relative to NOD.hIAPP 0/0 controls (Fig 2D, E). Consistent with reduced immune infiltration, CD45 (Ptprc) expression and T cell marker (Cd3d) expression was also reduced in whole islets (Fig 2C), as was the frequency of CD8 T cells in islets by flow cytometry (Fig 2F). No significant differences in CD4+, CD8+ (Fig 2G) or Treg (Fig 2H) frequency were observed in either spleen or the pancreatic lymph node between groups, implying a lack of a systemic perturbation in immune cell subsets. To further investigate whether NOD.hIAPP Tg/0 mice have a normal functioning systemic immune response, we immunized NOD.hIAPP Tg/0 and NOD.hIAPP 0/0 mice with Ova mRNA in a lipid nanoparticle vaccine by ip injection and measured the generation of anti-Ova antibodies. No difference in serum anti-Ova IgG1 concentrations were observed, indicative of an intact antigen presentation and immune response to systemically administered antigen (Fig 2I). Next, we examined whether splenocytes from pre-diabetic NOD.hIAPP Tg/0 mice could induce diabetes in immunocompromised NOD.Prkdc^Scid/Scid^ (i.e. NOD SCID) mice (Fig 2J). Splenocytes from NOD.hIAPP Tg/0 mice and NOD.hIAPP 0/0 mice induced diabetes at a similar rate, again indicating an intact autoimmune capacity of splenocytes from NOD.hIAPP Tg/0 (Fig 2K). Collectively, these data show that NOD.hIAPP Tg/0 are protected from immune infiltration, corresponding to reduced MHC Class II and APC activation in islets but normal systemic immune responses.

### NOD.hIAPP Tg/0 beta cells are dysfunctional but do not evade autoimmunity

To elucidate the mechanism of diabetes protection in hIAPP Tg/0 mice, we next examined the beta cell phenotypes of NOD.hIAPP Tg/0 mice prior to the onset of diabetes. In 8-week old mice, before the onset of diabetes, NOD.hIAPP Tg/0 female mice had impaired glucose tolerance in comparison to littermate controls (Fig. 3A, p=0.03) despite no differences in body weight (Fig 3B). A similar phenotype was observed in male NOD.hIAPP Tg/0 mice (Fig S1C-D). This mimics the well-established beta cell phenotype observed in hIAPP Tg/0 mice on non-autoimmune backgrounds. Despite no statistically significant differences in insulin levels following glucose stimulation, a trend toward higher insulin levels in vivo was apparent in female NOD.hIAPP Tg/0 vs. NOD.hIAPP 0/0 mice (Fig 3C). Ex vivo analysis revealed glucose-stimulated insulin secretion (Fig 3D) and insulin content (Fig 3E) to be higher from islets isolated from female NOD.hIAPP Tg/0 mice in comparison to islets isolated from and NOD.hIAPP 0/0 mice. This difference was abolished upon normalization to insulin content (Fig 3F). Male NOD.hIAPP Tg/0 mice also showed perturbations of in vivo and ex vivo stimulated-insulin secretion but had no difference in insulin content (Fig S1E-H). Histology analysis confirmed the development of islet amyloid in NOD.hIAPP Tg/0 female mice (Fig 3G). To determine whether hIAPP-induced perturbations in beta cells evade autoimmune attack, we generated immunocompromised NOD SCID.hIAPP Tg/0 and NOD SCID.hIAPP 0/0 mice and adoptively transferred splenocytes from diabetic NOD mice (Fig 3H). Diabetogenic splenocytes induced diabetes at a faster rate in NOD SCID.hIAPP Tg/0 recipients than in NOD SCID.hIAPP 0/0 controls (Fig 3I), demonstrating that beta cells expressing hIAPP do not evade autoimmune destruction. In summary, these data indicate that NOD.hIAPP Tg/0 mice develop islet amyloid and associated beta cell phenotypes similarly to other hIAPP Tg/0 strains and that these perturbations per se do not prevent autoimmune attack by diabetogenic T cells.

**Figure 3.**
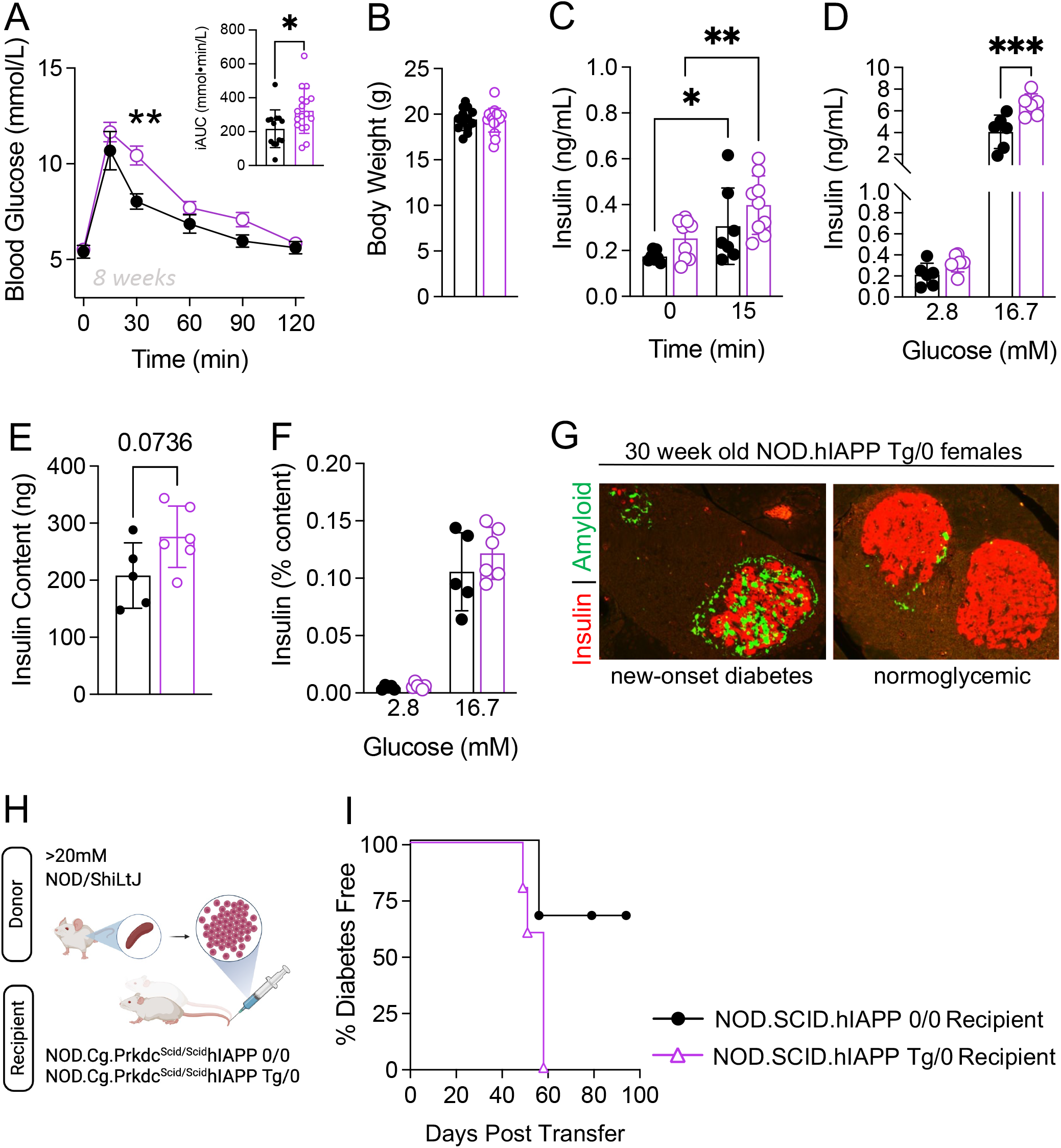
NOD.hIAPP Tg/0 beta cells are dysfunctional but do not evade autoimmunity. A) Glucose tolerance (1g/kg, i.p) of 8 week-old female NOD.hIAPP Tg/0 (n= 17) and control NOD.hIAPP 0/0 (n=13) mice. Data were analysed using a repeated measures mixed effects analysis. Individual incremental AUCs from experimental and control mice were analysed with the unpaired t-test. (B) Body weights at 8 week-old NOD.hIAPP Tg/0 (n= 17) and NOD.hIAPP 0/0 (n=13) mice. Data were analyzed using the Mann- Whitney test. (C) In vivo glucose-stimulated (1g/kg, i.p) plasma insulin levels from 8-week-old NOD.hIAPP Tg/0 (n= 9) and NOD.hIAPP 0/0 control (n=7) mice. Data were analyzed using a repeated measures mixed effects analysis. D) In vitro glucose-stimulated insulin secretion (GSIS) from islets isolated from 8-week-old NOD.hIAPP Tg/0 (n= 6) and control NOD.hIAPP 0/0 (n=6) mice. Data were analyzed using a mixed effects analysis E) Insulin content from islets isolated from 8-week-old NOD.hIAPP Tg/0 (n= 6) and control NOD.hIAPP 0/0 (n=5) mice. F) In vitro GSIS data normalized to insulin content from islets isolated from 8-week-old NOD.hIAPP Tg/0 (n= 6) and control NOD.hIAPP 0/0 (n=5) mice. G) Representative images of islet amyloid severity in newly-diabetic and normoglycemic 30 week old NOD.hIAPP female mice. H) Schematic overview of adoptive transfer of diabetic splenocytes from NOD donor mice into NOD SCID.hIAPP 0/0 or NOD SCID.hIAPP T/0 recipient mice (made in BioRender). I) Diabetes incidence curve of NOD SCID.hIAPP 0/0 (black solid circular symbol, n=3) or NOD SCID.hIAPP T/0 (pink open triangular symbol, n=5) recipient mice that have received splenocytes from diabetic NOD donor mice. Data were analyzed via logrank test. Data are shown as mean +/- SD. p < 0.05 (*), p < 0.01 (**),p < 0.0001 (****).

### hIAPP Tg/0 islets are not protected from alloimmune rejection

Finally, we examined whether the disruption of MHC Class II antigen presentation in islet macrophages may also delay immune attack of transplanted islets in an allogeneic islet transplant model. hIAPP Tg/0 and hIAPP 0/0 islets from F1 C57BL/6J x FVB/NJ donor mice were transplanted into STZ-induced diabetic Balb/cJ recipient mice and in direct contrast to the autoimmune models, allotransplant rejection of hIAPP Tg/0 islets was modestly accelerated relative to hIAPP 0/0 controls (Figure 4A-B). Notably, in allotransplant rejection, antigen recognition occurs through both APCs processing and presenting alloantigen and direct recognition of allogeneic MHC molecules on transplanted cells. This suggests that processing-dependent antigen presentation pathways are disrupted by IAPP aggregates whereas direct recognition of alloantigen is not.

**Figure 4.**
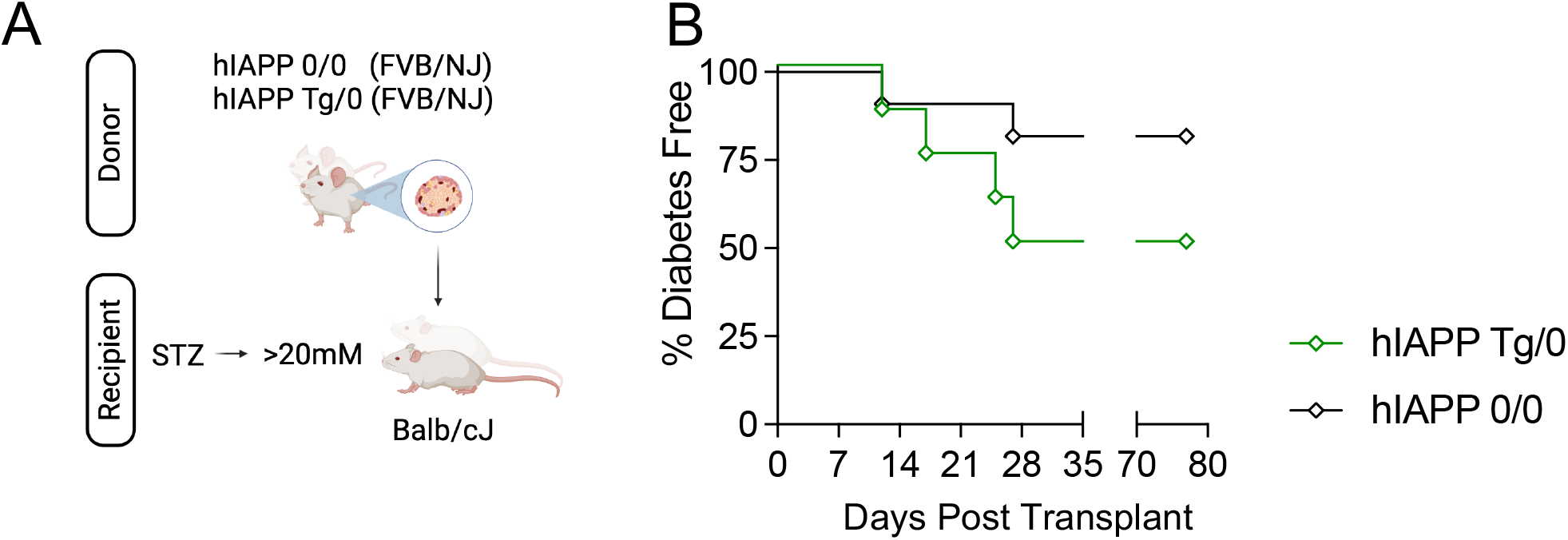
hIAPP expression does not delay islet allograft rejection. A) Schematic overview of islet transplantation studies (made in BioRender). In brief, isolated islets from 25 week-old hIAPP 0/0 or hIAPP Tg/0 male mice on the FVB/NJ background were transplanted under the kidney into STZ-induced diabetic 8-week male BALB-cJ mice. B) Diabetes incidence of male BALB-cJ mice with transplanted islets isolated from either hIAPP 0/0 control (black open diamonds, n=11) or hIAPP Tg/0 control (green open diamonds, n=8) mice. Data were analyzed via a logrank test.

### Phagocytosis of IAPP aggregates disrupt MHC Class II antigen presentation

To examine the mechanism of disrupted antigen presentation in hIAPP expressing mice, we modelled the direct effects of hIAPP on APCs by incubating BMDCs with either synthetic human or rodent IAPP in vitro. Synthetic hIAPP, in contrast to rIAPP, spontaneously forms IAPP aggregates in vitro, mimicking islet amyloid formation. We found that hIAPP pre-treatment inhibited the LPS-induced increase in MHCII^bright^ BMDCs (38.0±0.8 vehicle + LPS vs. 22.9±1.7% hIAPP + LPS; Fig 5A-B). This was not phenocopied by synthetic rIAPP (38.0±0.8% vs. 35.9±1.2% rIAPP + LPS), indicating the reduction in MHCII^bright^ BMDCs was not a function of monomeric IAPP but rather specific to IAPP aggregates.

**Figure 5.**
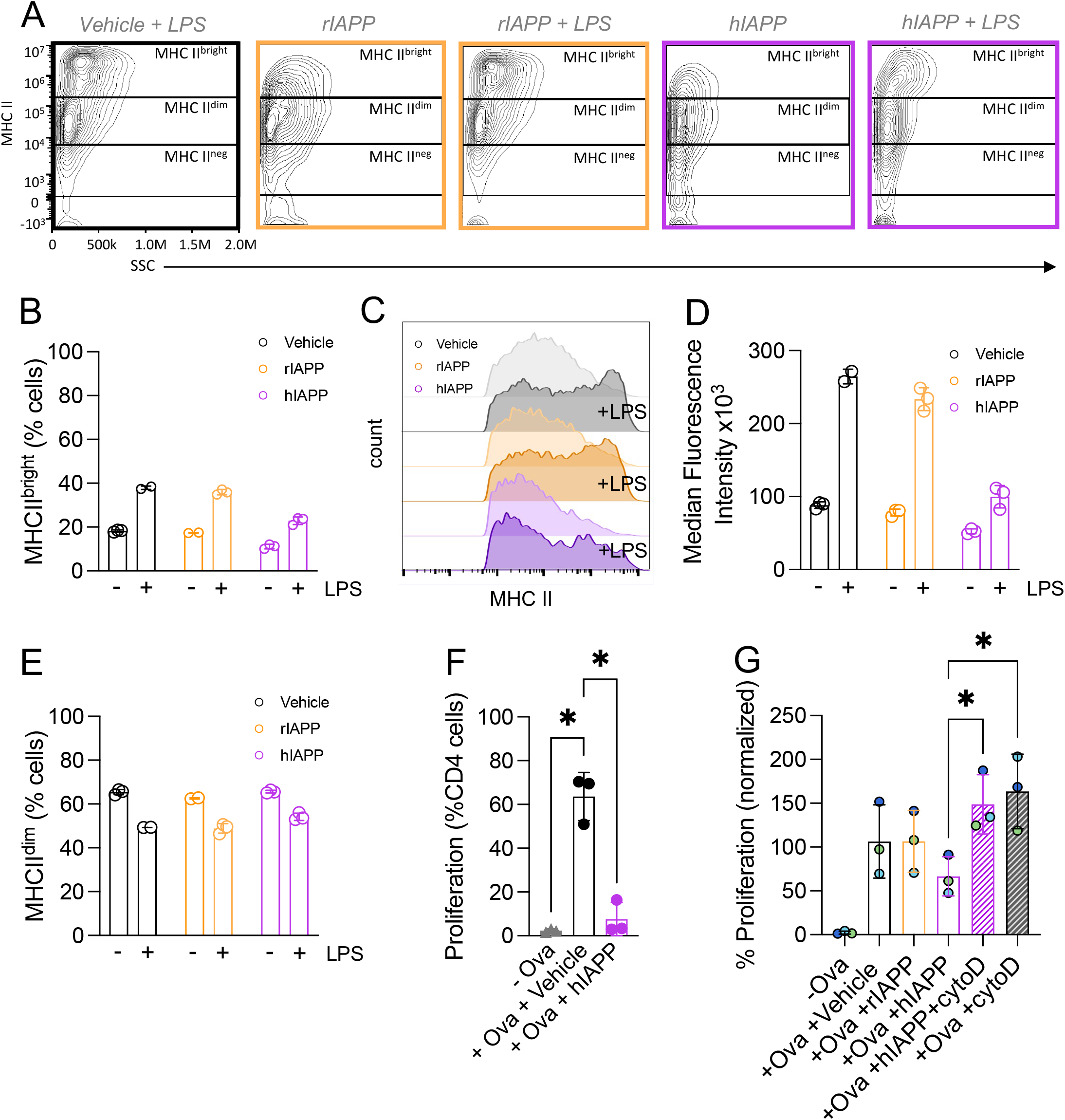
IAPP aggregates disrupt MHC Class II antigen presentation. A-E) Data from BMDCs pre-treated overnight with or without LPS (10ng/ml) in addition to vehicle (black), rIAPP (yellow) or hIAPP (pink) (10uM). A) Representative FACS plots showing MHCII levels in treated BMDCs. B) Frequency of treated BMDCs expressing high (MHCII^bright^) levels of MHCII. C) Histogram plot of treated BMDCs. D) Median Fluorescence Intensity quantification of D). E) Frequency of treated BMDCs expressing low (MHCII^dim^) levels of MHCII. F) Proliferation of Ova-specific CD4 T cells (% CD4 T cells) co-cultured with BMDCs pretreated with Ovalbulmin (Ova) and either vehicle (black) or hIAPP (pink). Data were analysed using a one-way ANOVA test. G) Proliferation of Ova-specific CD4 T cells co-cultured with BMDCs pretreated with Ovalbulmin (Ova) and either vehicle (black), rIAPP (yellow), hIAPP (pink, no pattern), hIAPP and cytoD (pink, with pattern) or cytoD only (black, with pattern). Data were normalized to the mean proliferation (% CD4 T cells) of Ova-specific CD4 T cells co-cultured with BMDCs pretreated with Ova only. Symbols of the same colour across different groups represent data generated within the same experiment. Data were analysed using two-way ANOVA. Data are shown as mean +/- SD. p < 0.05 (*), p < 0.01 (**),p < 0.0001 (****).

Similarly, hIAPP prevented the LPS-induced increase in the mean fluorescence intensity of MHCII on BMDCs (Fig 5C, D). No differences in MHCII^dim^ BMDCs were observed between groups (Fig 5E). To examine whether this decrease in MHC Class II led to a functional difference in antigen presentation, we examined the effect of hIAPP on the ability of BMDCs to present the model antigen Ova to Ova-specific CD4+ T cells. Relative to control BMDCs, when hIAPP pre-treated BMDCs were pulsed with Ova peptide and then co-cultured with Ova-specific CD4+ T cells, antigen-specific T cell proliferation was dramatically reduced (Fig 5F). As phagocytosis of IAPP aggregates has previously been implicated in its pro-inflammatory effects on macrophages [17, 18], we set out to examine whether hIAPP-impaired antigen presentation was dependent on phagocytosis of hIAPP aggregates. BMDCs were co-treated with both hIAPP and cytochalasin D, a reversible phagocytosis inhibitor, to prevent phagocytosis of hIAPP aggregates. Subsequently, BMDCs were washed and pulsed with Ova peptide antigen, then washed again and co-cultured with Ova-specific T cells. Phagocytosis inhibition attenuated the inhibitory effects of hIAPP treatment on antigen presentation and normal levels of Ova-specific CD4^+^ T cell proliferation resulted (Fig 5G). Collectively these results show that direct phagocytosis of IAPP aggregates by APCs disrupts MHC Class II antigen presentation and activation of antigen-specific CD4+ T cells.

## Discussion

MHC Class II antigen presentation is the most down-regulated pathway in islet macrophages during the early stage of islet amyloid formation in hIAPP Tg/0 mice. Consequently, we generated novel strains of mice carrying amyloidogenic IAPP on the NOD genetic background to probe the effect of islet amyloid in autoimmune diabetes. NOD mice expressing hIAPP Tg/0 developed glucose intolerance at an early age and had demonstrable insulin secretion alterations and islet amyloid in islets, consistent with phenotypes of hIAPP Tg/0 mice. Despite these impairments in beta cell function, autoimmune diabetes was significantly delayed in NOD mice expressing hIAPP by either transgene or knock-in to the endogenous mouse Iapp locus. This corresponded to less T cell infiltration and a corresponding reduction in MHC Class II and markers of APC activation in islets, implicating islet amyloid in disrupting MHC Class II antigen presentation and slowing autoimmune progression of diabetes. Splenocytes from hIAPP Tg/0 NOD mice effectively induced diabetes in immune compromised mice and hIAPP Tg/0 NOD mice responded normally to Ova immunization, indicating the delayed diabetes is not driven by impaired autoreactive T cells and that impaired antigen presentation is likely localized to pancreatic APCs. We examined the direct effects of IAPP aggregates on APCs in vitro, and confirmed that IAPP aggregates disrupted antigen presentation directly in APCs through phagocytosis.

The findings of this paper are strengthened using multiple experimental models to triangulate the consistent effect of amyloidogenic IAPP on APCs. An impact on MHC Class II antigen presentation was initially identified in an unbiased scRNAseq approach, followed by confirmation of functional reductions in antigen presentation in vitro and confirmation of alterations in antigen presentation culminating in protection from autoimmune diabetes in two independent strains of NOD mice expressing hIAPP. Importantly, this controls for artifacts that could result from overexpression of the hIAPP transgene in beta cells (or other cells for that matter), and also for the proximity of any Idd genes around the hIAPP transgene insertion site (chromosome 15) or Iapp locus (chromosome 6, Iapp also being an Idd locus itself and important for the formation of hybrid insulin peptide antigen, HIP6.9) [42–45]. We also confirmed that the effect of hIAPP on MHC Class II antigen presentation was a direct effect on APCs through IAPP aggregates specifically, as non-aggregating rodent IAPP did not impact antigen presentation or MHC Class II surface expression. This is consistent with previous reports that Iapp knockout mice do not have altered diabetes progression [44]. This is an important point considering evidence that IAPP itself may be an autoantigen and its ability to form hybrid insulin peptide antigens recognized by CD4 T cells in NOD mice [42–45]. In addition, we confirmed that the delay in diabetes caused by hIAPP expression in NOD mice was not due to decreased systemic antigen presentation (normal response to Ova immunization) or T cell function (as NOD.hIAPP Tg/0 splenocytes can adoptively transfer diabetes), or altered beta cell function (NOD SCID.hIAPP Tg/0 beta cells do not evade autoimmunity following adoptive transfer of diabetogenic T cells). A limitation of this study is that we did not confirm our findings on impaired antigen presentation in a human model system (e.g. human islets or APCs with HLA-matched human T cells). These studies would be of value in determining whether our observation in mice may generalize to human type 1 diabetes.

The protective effect of hIAPP in the NOD mouse model of type 1 diabetes was somewhat surprising given the extensive evidence that IAPP aggregates and islet amyloid induce islet inflammation, impair beta cell function and contribute to beta cell failure and loss in type 2 diabetes (reviewed in [46]). IAPP aggregates interact directly with macrophages through TLR2 and phagocytosis and induce proinflammatory cytokine secretion which contributes to beta cell dysfunction [15, 17, 18]. Indeed, we recently showed that targeting IAPP aggregates specifically with a monoclonal antibody improves the survival and function of human islet transplants in mice, again highlighting the deleterious effects of islet amyloid on beta cell function [47]. The deleterious impacts of amyloid on beta cell function are still present in both the presence and absence of autoimmunity, confirmed here by glucose intolerance in both hIAPP Tg/0 C57BL/6J x FVB/NJ mice and pre-diabetic NOD.hIAPP Tg/0 mice). The impact on islet inflammation was also still apparent, evident in the increased CD45+ cells in hIAPP Tg/0 C57BL/6J x FVB/NJ islets compared to littermate controls. Despite this, our findings suggest that in the presence of beta cell autoimmunity, the impact of IAPP aggregates on antigen presentation outweigh their direct induction of inflammation, as immune infiltration was decreased in NOD.hIAPP Tg/0 mice relative to controls. Also of interest is that rejection of hIAPP Tg/0 islets was accelerated in an allotransplant model, consistent with increased islet inflammation being present and further suggesting that IAPP aggregates impair antigen processing pathways but not direct recognition of alloantigens.

The mechanism by which IAPP aggregates impair MHC Class II antigen presentation appears to share striking similarities with the mechanism by which IAPP aggregates induce inflammation, particularly the induction of IL-1B secretion. Early IAPP aggregates interact through TLR2 to prime expression of proinflammatory cytokines, and phagocytosis of more mature IAPP aggregates leads to lysosomal disruption and consequent activation of the NLRP3 inflammasome, essential for maturation and secretion of IL-1B [15, 17, 18]. Here we show that disruption of MHC Class II antigen presentation is also dependent on phagocytosis of IAPP aggregates. Since MHC Class II antigen processing and loading occurs throughout the phagolysosomal system, we speculate that the known disruption of lysosomes by IAPP aggregates is also what disrupts MHC Class II antigen processing, leading to less MHC Class II antigen presentation on the cell surface. Whether this would also lead to the observed decrease in Ciita and MHC Class II presentation genes which we observed is unclear, but it is possible that a feedback loop could be induced by lysosomal disruption.

Based on our findings, we propose the following model to account for the protective effect of IAPP aggregates in autoimmune diabetes in the face of extensive evidence, including our own, that IAPP aggregates elicit islet inflammation and beta cell loss in type 2 diabetes. Islet macrophages are the main APCs resident in islets and directly interact with IAPP aggregates. As islet macrophages interact with IAPP aggregates, particularly through their phagocytosis, it elicits two opposing effects on the macrophage: 1) induction of proinflammatory cytokine secretion which promotes islet inflammation and beta cell dysfunction and 2) disruption of MHC Class II antigen presentation which reduces induction of adaptive immune responses. The outcome of these opposing effects on immunity, in regard to beta-cell protection or loss, depends on whether this occurs in the presence of beta cell autoimmunity. When autoimmunity is present, the protective effect of disrupted antigen presentation outweighs the induction of inflammation, resulting in protection. In the absence of autoimmunity, as in models of type 2 diabetes and syngeneic islet transplants, disrupted antigen presentation is inconsequential and the islet inflammation contributes to beta cell dysfunction. Similarly, in allogeneic islet transplants where multiple pathways to alloantigen recognition exist, disruption of MHC Class II antigen presentation does not impart a protective effect and the deleterious impact of islet amyloid on beta cell function is apparent.

What is the consequence and role of islet amyloid, recently found to be present in type 1 diabetes? Our data suggest that as IAPP aggregates form, perhaps in response to early beta cell dysfunction and beta cell death, islet amyloid may quell the pace of further autoimmune stimulation, though notably is not likely to impact the action of autoreactive CD8 T cells in recognizing MHC Class I antigens on islet cells. It is intriguing, far-fetched as it may be, to ponder connections between the ability of human but not mouse islets to naturally form islet amyloid, and the disparity between insulitis in humans and mouse insulitis, where even active insulitis in type 1 diabetes exhibits few immune cells per islet compared to the extensive infiltration of islets in NOD mice. Nevertheless, more studies are needed to clearly define the role of islet amyloid in type 1 diabetes.

## Acknowledgements

hIAPP knock-in mice were provided in kind from Dr. Steven Kahn (University of Washington Diabetes Institute, USA). Lipid nanoparticles were provided in kind by Integrated Nanotherapeutics.

RNAsequencing was performed by the BRC-Seq Next Gen Sequencing Core (University of British Columbia, Canada). The authors wish to thank Lisa Xu for providing her expertise and assistance with flow cytometry and FACS. The authors also wish to thank Søs Skovsø, PhD from Valkyrie Life Sciences who facilitated manuscript drafting and editing as well as data analysis, figure and schematic overview generation.

## Data Availability

RNAseq data will be deposited in Gene Expression Omnibus database upon publication. Any data that support the findings of this study are available from the corresponding author with reasonable request.

## Author’s relationships and activities

HD, SC and BV are cofounders of Integrated Nanotherapeutics. HD took on employment at Integrated Nanotherapeutics after her postdoctoral fellowship and completion of the experiments in this manuscript.

## Contribution statement

HD conceived, designed and conducted the research in this manuscript, analyzed the majority of data and wrote, edited and reviewed the manuscript. LS and IS helped with mouse monitoring.

TN conducted proliferation assays and BMDC experiments.

DN, MK assisted with scRNAseq experiment and JV conducted scRNAseq analysis.

SC designed and formulated LNPs for Ova immunization and INT provided these LNPs in kind. BV reviewed and edited the manuscript and is the guarantor of the data.

---

**Supplemental Figure 1.**
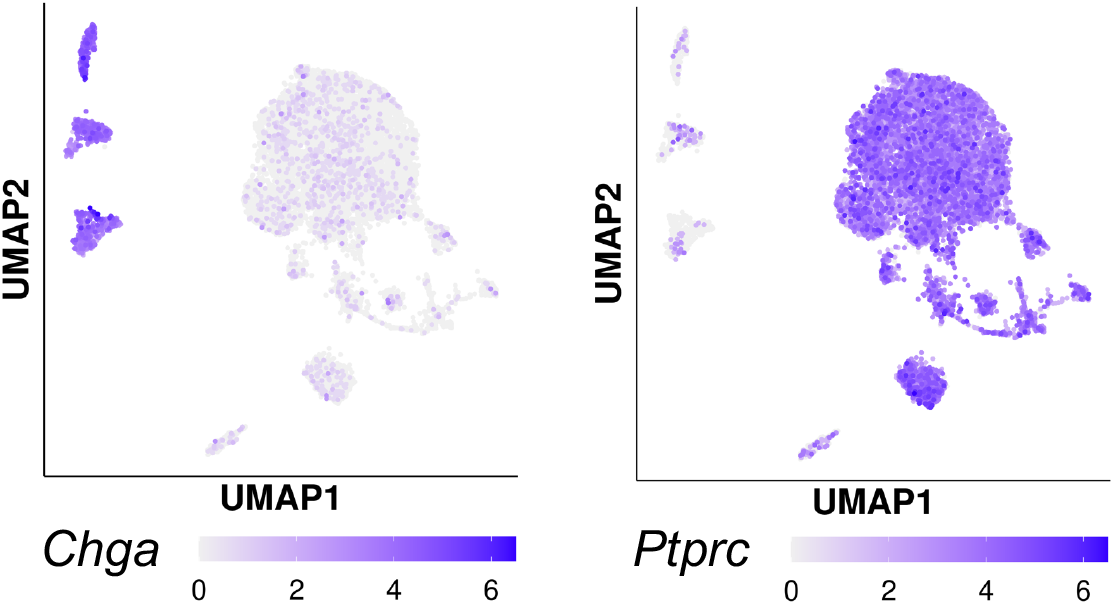
Expression of endocrine and immune markers in islet cell populations. A) Feature Plots showing Chromogranin A (ChgA) in endocrine cells and Protein Tyrosine Phosphatase Receptor Type C (Ptprc, also known as CD45) expression in immune cells.

**Supplemental Figure 2.**
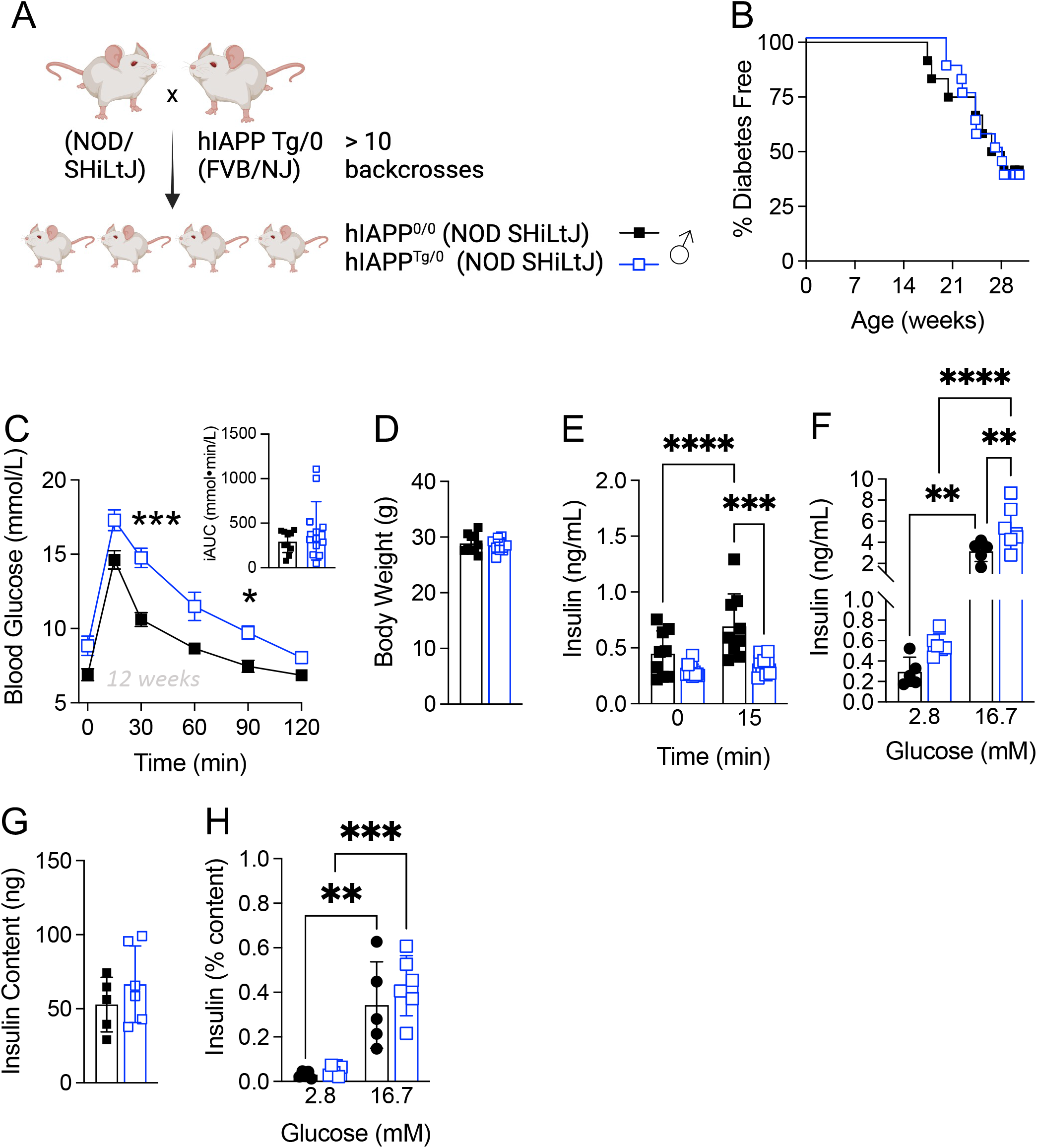
NOD.hIAPP Tg/0 male mice have the expected islet amyloid-induced beta cell phenotype. A-G) Data from male control (black, solid square) and hIAPP Tg/0 (blue, open square) transgenic mice on a NOD SHiLtJ background. A) Overview of hIAPP transgenic NOD/SHiLtJ animal model breeding strategy (made in BioRender) and diabetes incidence. hIAPP Tg/0 mice were fully backcrossed from an FVB/NJ background to a NOD SHiLtJ background (> 10 backcrosses). B) Diabetes incidence of male NOD.hIAPP Tg/0 (n= 13) and NOD.hIAPP 0/0 control (n=12) mice. Data were analyzed via a logrank test. C) Glucose tolerance (dose, i.p) of 12-week-old male NOD.hIAPP Tg/0 (n= 12) and control NOD.hIAPP 0/0 (n=10) mice. Data were analysed using a repeated measures mixed effects analysis. Individual incremental AUCs from experimental and control mice were analysed with the unpaired t-test. (D) Body weights at 12-week-old hIAPP Tg/0 (n= 10) and hIAPP 0/0 (n=10) mice. Data were analyzed using the Mann-Whitney test. (E) In vivo glucose-stimulated (1g/kg, i.p) plasma insulin levels from 12-week-old hIAPP Tg/0 (n= 9) and hIAPP 0/0 control (n=7) mice. Data were analyzed using a repeated measures mixed effects analysis. F) In vitro glucose-stimulated insulin secretion (GSIS) from islets isolated from 12-week-old hIAPP Tg/0 (n= 10) and control hIAPP 0/0 (n=9) mice. Data were analyzed using a mixed effects analysis G) Insulin content from islets isolated from 12-week-old hIAPP Tg/0 (n= 6) and control hIAPP 0/0 (n=5) mice. H) In vitro GSIS data normalized to insulin content from islets isolated hIAPP Tg/0 (n= 6) and control hIAPP 0/0 (n=5) mice. Data are shown as mean +/- SD. p < 0.05 (*), p < 0.01 (**),p < 0.0001 (****).

